# PFAS Compounds PFOA and Gen X are Teratogenic to Sea Urchin Embryos

**DOI:** 10.1101/2024.11.21.624751

**Authors:** Alexandra T. Lion, Sophie M. Bodine, Kelley R. McCutcheon, Mayank Ghogale, Santhan Chandragiri, Deema Abayawardena, Bikram D. Shrestha, Abigail Descoteaux, Kathryn Alvarez, J’nesse A. Balkman, Breelyn Cocke, Athula H. Wikramanayake, Jennifer Schlezinger, Joyce Y. Wong, Vivek N. Prakash, Cynthia A. Bradham

## Abstract

Per-and polyfluorinated substances (PFAS) are synthetic chemicals that are used to make fluoropolymer coatings found in many products, such as non-stick pans, clothing, cosmetics, and food packaging. These highly persistent molecules are known as “forever chemicals” since they neither degrade environmentally nor break down enzymatically within biological systems. PFAS compounds readily contaminate water sources, and as a result, certain PFAS molecules have bioaccumulated in exposed species including humans. The purpose of this study was to define the effect of two PFAS molecules, the ostensibly more toxic perfluorooctanoic acid (PFOA) and the more recent, reportedly safer chemical hexafluoropropylene oxide dimer acid (Gen X), on the development of *Lytechinus variegatus* sea urchin embryos. We examined the effects of PFOA and Gen X on development and patterning using morphological analysis, immunostaining, HCR-FISH, and Particle Image Velocimetry (PIV). The results show that both PFAS compounds are teratogenic to sea urchin embryos. PFOA and Gen X each function at different intervals during development and provoke distinct phenotypic and gene expression outcomes. Despite beliefs that Gen X would be a safer alternative, our findings indicate that Gen X has earlier and more severe effects on endomesoderm and dorsal-ventral axis specification, neural development and function, and pattern formation compared to PFOA. These results illustrate the dangerous teratogenic potential of environmentally accumulating PFAS like Gen X, underscoring the negative ecological implications that accompany continuing commercial and industrial use of PFAS in the absence of remediation strategies.

**HIGHLIGHTS:**

- Gen X is more teratogenic in sea urchins than PFOA
- Gen X acts earlier than PFOA and perturbs axial and germ layer specification
- Each PFAS reduces neural numbers and perturbs ciliary behavior and swimming

## Introduction

Per-and polyfluroalkyl substances (PFAS) are synthetic chemicals that are used to make fluoropolymer coatings found in many products, such as non-stick pans, clothing, cosmetics, and food packaging; PFAS are also used in aqueous, film-forming foam used in firefighting (Ackerman Grunfeld et al., 2024; Buck et al., 2011; Ding et al., 2020; Gaballah et al., 2020; Gebreab et al., 2020; Landrigan et al., 2020; Wang et al., 2017). Known colloquially as “forever chemicals,” PFAS are highly stable and persistent in the environment due to the chemical stability of the C-F bond and their lack of both natural and enzymatic degradation. These characteristics have allowed some PFAS to bioaccumulate in exposed species (Kwiatkowski et al., 2020; Macorps et al., 2022; Pan et al., 2021; Son et al., 2020). Since many directly exposed animals, i.e. in marine environments, are low on the food chain, PFAS exhibit broad bioaccumulation across many species including humans.

Humans are also exposed to PFAS through a variety of pathways in addition to the consumption of animal products, including via drinking water, the air, food packaging materials, and cosmetics (Blake & Fenton, 2020; Cousins et al., 2021; Davidsen et al., 2021). Nearly all humans have a detectable percentage of PFAS in their bloodstream (Davidsen et al., 2021; Ding et al., 2020; Gebreab et al., 2020). PFAS are associated with detrimental health effects including suppressing immune responses to vaccines, liver toxicity, increasing serum cholesterol, increasing the risk of pregnancy-induced hypertension, and increased risk of kidney and testicular cancer (Agency for Toxic Substances and Disease Registry, 2021; Blake & Fenton, 2020; Fenton et al., 2017; Kotlarz et al., 2020; Park et al., 2024; Phelps et al., 2023; Rericha et al., 2023). PFAS exposure has negative effects on development, leading to morphological defects, neurological defects and neurotoxicity, and low birthweights (Carstens et al., 2023; Fan et al., 2023; Mahapatra et al., 2023; Rericha et al., 2023). Moreover, PFAS accumulate in the human placenta, allowing them to be passed to the fetus *in utero* and promoting high developmental exposure levels (Blake et al., 2020; Chowdhury et al., 2024; Kilari et al., 2024; Lemos et al., 2024; Mamsen et al., 2019). This observation in particular calls for extensive study of the impacts of PFAS on embryonic development.

This study focuses on two PFAS, perfluorooctanoic acid (PFOA), and hexafluoropropylene oxide dimer acid (HFPO-DA) fluoride, known as Gen X. PFOA has been linked to toxic effects in humans and model organisms and is a teratogen (Fenton et al., 2017; Gaballah et al., 2020; Kotlarz et al., 2020; Liu & Irudayaraj, 2020; Yu et al., 2021). As such, PFOA has been generally phased out of production in the United States and replaced with Gen X, based on initial studies indicating that Gen X bioaccumulates less than PFOA. Despite its increased use, the biological impacts of Gen X have been understudied compared to PFOA and other “legacy” PFAS (Barragan et al., 2023; Davidsen et al., 2021; Gebbink et al., 2017; Gebbink & van Leeuwen, 2020; Phelps et al., 2023). Despite its reportedly shorter half-life, studies have indicated that the toxic effects of Gen X are similar to those of PFOA (Blake et al., 2020; Fenton et al., 2017; Gebbink & van Leeuwen, 2020; Gomis et al., 2018). Both PFOA and Gen X impaired human cardiomyocyte differentiation, suggesting Gen X may also be a developmental toxicant (Davidsen et al., 2021). In addition, when differences in toxokinetics were considered, Gen X appeared to have higher toxic potency in serum and liver than PFOA (Blake et al., 2022; Gomis et al., 2018; Robarts et al., 2024; Zhang et al., 2024).

Thus, further studies of the biological effects of Gen X on developing embryos are warranted, because Gen X may not be the “safer” replacement it was initially thought to be. It is also important to understand the adverse ecosystem effects of environmental PFAS contamination. Run-off from factories, production plants, airports, and military bases have caused wide-spread PFAS contamination of fresh waters; PFAS have also entered the oceans in large quantities. Despite this reality, little is known about the consequences of PFAS accumulation in marine species (Hayman et al., 2021; Khan et al., 2023; Landrigan et al., 2020; Lukić Bilela et al., 2023).

Here, we utilize *Lytechinus variegatus* (Lv) embryos as a sentinel marine model organism to study the impacts of PFOA and Gen X. *L. variegatus*, commonly known as the green sea urchin, is an excellent model organism for studying effects on development. *L. variegatus* embryos are transparent and morphologically simple, making their development accessible. Development and embryogenesis are rapid, with larval development well-underway 48 hours post-fertilization (hpf) (Adonin et al., 2021). The surprisingly high genetic similarly of sea urchins to vertebrates, including humans, makes them a suitable model for genetic analysis (Adonin et al., 2021; Bradham et al., 2006; Davidson, 2006; McClay, 2011; Sodergren et al., 2007). Their sensitivity to environmental changes and pollutants makes sea urchin embryos, including *L. variegatus*, excellent sentinel species (Arnberg et al., 2018; Cossa et al., 2024; Dorey et al., 2022; Duvane & Dupont, 2024; Moreno-García et al., 2022; Pinsino & Matranga, 2015; Przeslawski et al., 2015; Sarly et al., 2023; Sato et al., 2018) that predict biological consequences for other marine animals that exhibit less sensitivity. The goal of this study was to define the effects of PFOA and Gen X on development and patterning in *L. variegatus* embryos, and to elucidate how PFAS exposure perturbs these processes.

## Results

### PFOA and Gen X have effects on secondary mesenchyme cell (SMC) and endodermal derived structures

To determine the effect of exposure to PFOA or Gen X (Fig. 1A1-2) on the development of sea urchin embryos, *Lytechinus variegatus* embryos were treated with either PFOA or Gen X at fertilization. Compared to time-matched controls (Fig. 1B1-3), larvae treated with PFOA appeared mildly stunted (Fig. 1C1-3), while Gen X treated embryos exhibit a severe delay to gut development and dramatically perturbed morphology (Fig. 1D1-3), along with a notable absence of secondary mesenchyme cell (SMC)-derived pigment cells at 48 hours post-fertilization (hpf) (Fig. 1D3). The doses required to obtain these phenotypes are relatively high, and this likely reflects the hydrophilicity of each molecule, presenting a barrier to diffusive entry into the embryo that is likely overcome only at these concentrations.

**Figure 1:**
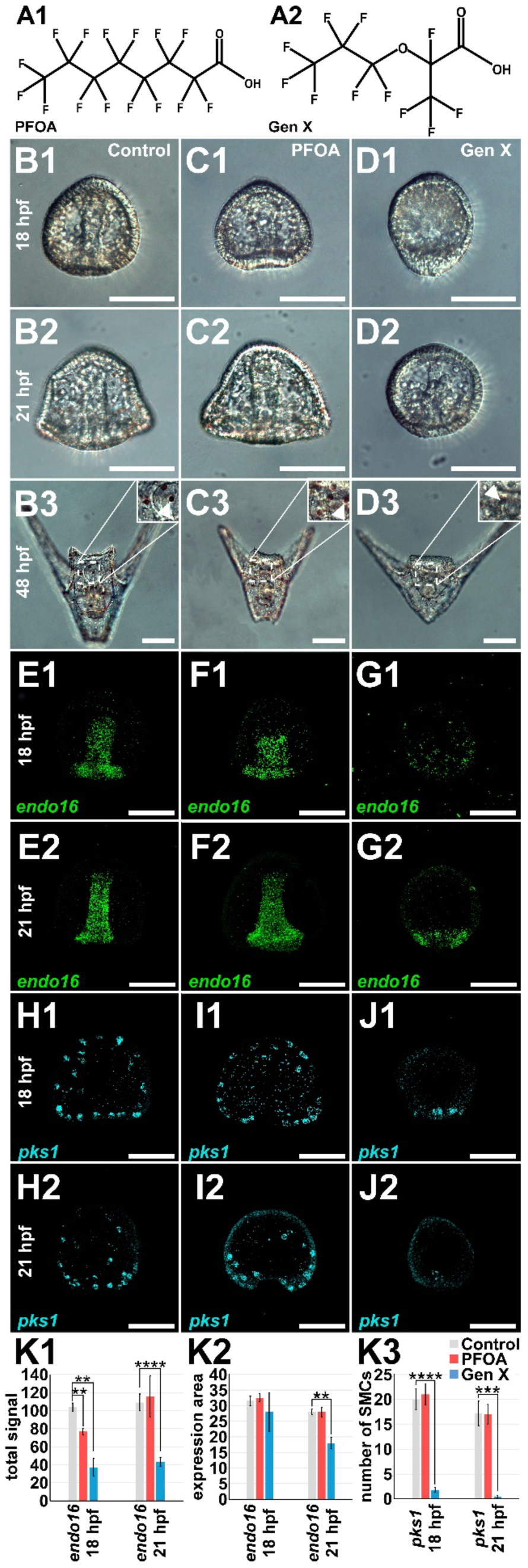
Gen X is sufficient to perturb SMC and endodermal patterning. **A.** PFAS molecular structures are shown for perflurooctanoic acid (PFOA, C_8_HF_15_O_2_) (A1) and hexafluoropropylene oxide dimer acid (Gen X, C_6_HF_11_O_3_) (A2). **B-D.** Phenotypes for control (B), PFOA (300 μM, C), or Gen X (250 μM, D). **E-J.** Control and PFAS-treated embryos were subject to HCR-FISH for Lv*endo16* (E-G) or Lv*pks1* (H-J) at 18 (1) and 21 (2) hpf. **K.** The total expression level (1) and normalized area (2) for endo16, and the number of pks1-positive SMCs (3) are plotted as the average ± s.e.m. ** p< 0.005, *** p< 0.0005, and **** p < 0.00005; otherwise, insignificant (*t*-tests). Scale bars are 50 microns here and in all other figures unless indicated.

To better assess endoderm and SMC perturbation with PFAS exposure, we performed HCR-FISH for *endo16* and *pks1*, which are expressed in the gut and pigment cells, respectively (Calestani et al., 2003; Calestani & Rogers, 2010; Romano & Wray, 2006). In PFOA-treated embryos, *endo16* expression is significantly reduced at 18 hpf, but recovers by 21 hpf (Fig. 1F1-2, K1). Gen X-treated embryos show a significant and dramatic reduction in *endo16* expression at both 18 and 21 hpf, as well as a significant reduction in the *endo16* expression area at 21 hpf (Fig. 1G1-2, K1-2). Gen X but not PFOA exposure significantly and dramatically inhibited SMC specification, with a significant reduction in SMC numbers per embryo at both 18 and 21 hpf (Fig. 1H1-J2, K3). Thus, PFAS exposure perturbs endomesoderm specification in sea urchin embryos, with Gen X exhibiting more severe and long-lasting impacts on both tissues.

### PFOA and Gen X perturb sea urchin skeletal patterning and PMC migration

Along with SMCs, the embryo also contains primary mesenchyme cells (PMCs) that initially serve as the major organizer, then later differentiate as skeletogenic mesenchyme cells (Ettensohn & Adomako-Ankomah, 2019; Gustafson & Wolpert, 1961a; Gustafson & Wolpert, 1961b; Horstadius, 1939; Lyons et al., 2014; Wolpert & Gustafson, 1961). The PMCs adopt spatial positions within the blastocoel in response to ectodermal cues whose spatial expression dictates the skeletal pattern (Armstrong et al., 1993; Duloquin et al., 2007; Malinda & Ettensohn, 1994; Piacentino, Zuch, et al., 2016). To determine the effects of PFOA or Gen X exposure on larval skeletal development, we visualized the skeletal morphology in 48 hpf larvae that were treated with either chemical at fertilization (Fig. 2A-D). PFOA-treated embryos exhibited significant numbers of secondary anterior skeletal losses and rotations around the anterior-posterior (AP) axis, while Gen X-treated embryos show mainly secondary element duplications, rotations around the dorsal-ventral (DV) axis, and spurious elements (Fig. 2A-D, S1). Transformed elements arose more frequently and significantly with PFOA exposure compared to Gen X, although the overall number of embryos exhibiting transformations was relatively low in both cases (Fig. S1B). Thus, exposure to PFAS compounds PFOA or Gen X is sufficient to elicit skeletal patterning defects in sea urchin embryos, with each treatment producing distinct defects.

**Figure 2:**
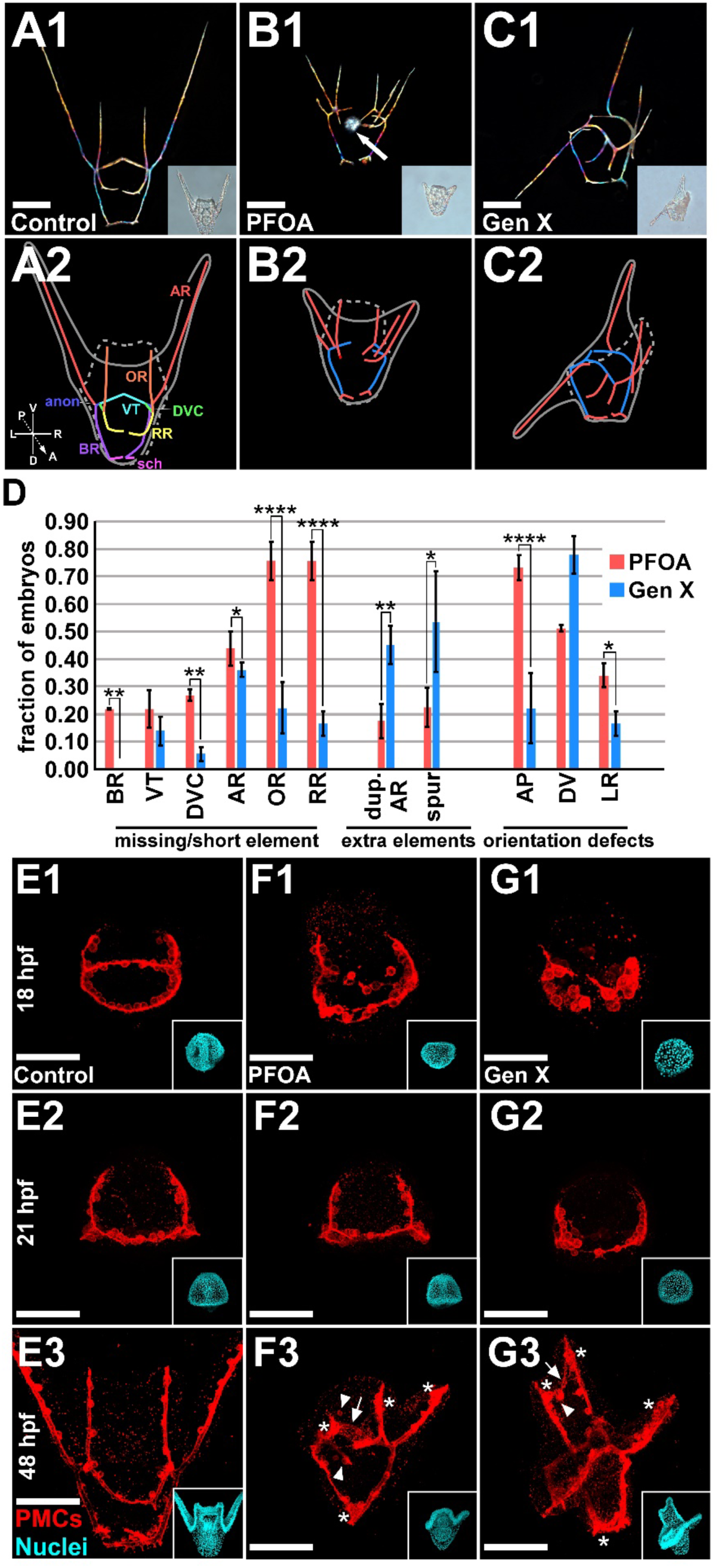
PFOA and Gen X are sufficient to perturb skeletal patterning and PMC migration. **A-C** The skeletons within control (A), PFOA-(300 μM, B), and Gen X-treated (250 μM, C) larvae are shown at 48 hpf as live birefringence images with the corresponding DIC images inset (1) and as schematics (2) with primary skeletal elements in blue and secondary skeletal elements in red. Non-skeletal birefringence signals in the gut (arrow in B1) appear to reflect ingested PFOA. Skeletal rods that are indicated in A2 include VT ventral transverse rods; DVC dorsal-ventral connecting rods; anon anonymous rods; BR body rods; AR aboral rods; OR oral rods; and RR recurrent rods. **D**. Skeletal defects in PFOA- and Gen X-treated embryos at 48 hpf are plotted as averages ± s.e.m. * p < 0.05; ** p < 0.005; **** p < 0.00005; otherwise, insignificant (*t*-tests); n=36-41 per condition. **E-G.** PMCs were immunolabeled in control embryos (E) and in embryos treated with PFOA (F) or Gen X (G) at 18 (1), 21 (2) or 48 (3) hpf; corresponding Hoechst-stained images are inset. Arrows, arrowheads, and asterisks in F3 and G3 indicate abnormal filopodia, ectopically positioned PMCs, and abnormally large PMC clusters, respectively. See also Fig. S1-3.

In addition to the birefringence of their skeletons, some PFAS-treated embryos also exhibited a diffuse birefringence signal within their guts (Fig. 2B1, arrow). To test whether that signal reflects ectopic biomineralization, we employed calcium-binding fluorophores to label the skeleton (Descoteaux et al., 2023). Those results show that the skeleton but not the gut-specific birefringence labels with these fluorophores (Fig. S2), indicating that the birefringence in the gut is not due to local calcium accumulation. We therefore conclude that the signal within the gut reflects ingested PFOA, which is naturally birefringent (Butnor et al., 2020). The polychrome labeling experiments also show that the skeleton develops with normal timing in PFOA-treated embryos. During skeletal patterning and biomineralization, the PMCs extend thin filopodia that are thought to sense patterning cues expressed by the ectoderm; those cues direct PMC migration and positioning into a stereotypical spatial pattern comprised of two ventrolateral clusters of PMCs that are connected by chains of cells which form a posterior ring that encircles the blastopore and hindgut, along with two cords of cells that extend anteriorly from ventrolateral clusters (Armstrong et al., 1993; Ettensohn & Malinda, 1993; Miller et al., 1995; Piacentino et al., 2015; Piacentino, Zuch, et al., 2016). The PMCs also extend thick cables which join to form a tubular syncytium that connects the cells, and into which the skeleton is secreted during biomineralization (Kahil et al., 2020; Khor et al., 2023).

To determine whether the spatial positioning of the PMCs is perturbed by PFAS exposure, PMCs were visualized using immunolabeling (Fig. 2E-G). PFOA- and Gen X-treated embryos each display abnormally positioned PMCs at 18 hpf, along with abnormal filopodia, abnormal PMC clusters, and ectopically positioned PMCs. While both PFOA and Gen X elicit a morphological delay, these delays are more pronounced for Gen X, whose PMCs are delayed in both the formation of the initial ring-and-cords pattern, and the progression from it (Fig. 2G, S3). These findings show that PFOA and Gen X treatment each result in abnormal migration and spatial organization of the PMCs; the stronger perturbations evident in Gen X-treated embryos are consistent with the more pronounced skeletal patterning defects that it elicits compared to PFOA.

### PFOA and Gen X exert their effects during defined temporal windows

To determine when PFOA and Gen X perturb *L. variegatus* development, timed PFOA or Gen X (Fig. 3A-D) additions and removals were performed at a range of developmentally relevant time points. After reaching the larval stage (48 hpf), the resulting morphology was then scored, using skeletal patterning as a read-out.

**Figure 3.**
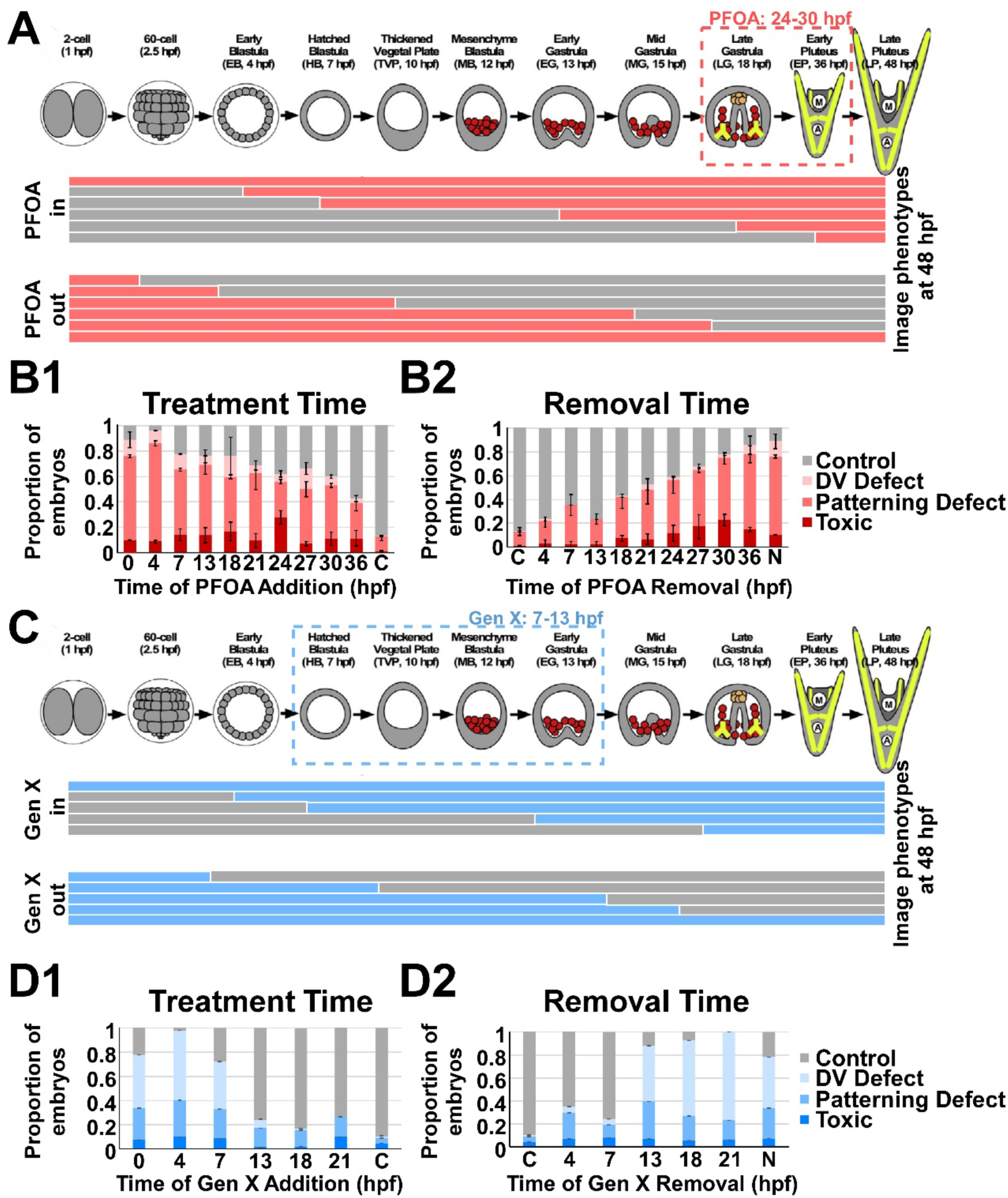
PFOA and Gen X treatment exert their effects during defined and distinct temporal windows. **A, C.** The schematic depicts the variable PFAS exposure windows for different addition (upper bars) or removal times (lower bars) for PFOA (A) or Gen X (C). The developmental time course schematic was adapted from Hogan et al, 2020. **B, D.** Embryos were treated with PFOA (300 μM, B) or with Gen X (250 μM, D) at the indicated time points through development (1) or treated at fertilization, then washed out of the PFAS at the indicated time points (2). The resulting embryos were scored at 48 hpf as indicated; the results are plotted as averages ± s.e.m. The temporal windows of action for PFOA and Gen X are shown as dotted boxes in A and C.

The dorsal-ventral axis is specified before and during the installation of skeletal patterning cues (Bradham & McClay, 2006; Hardin et al., 1992; Hardin & Armstrong, 1997; Hogan et al., 2020)). Loss of DV specification results in radialized skeletal patterns due to broad disruption of ectodermal specification. DV defects that are more mild lead to other distinct skeletal patterns, including partial ventral expansion as indicated by a very wide angle between the aboral rods and/or the presence of supernumerary aboral rods (AR) extending from the ventral-transverse rods (VT). In contrast, bona fide patterning defects are morphologically distinct (see Fig. 2) and do not arise from underlying DV aberrations.

The skeletal patterns obtained in these temporal experiments were scored for DV defects versus patterning defects. We found that PFOA perturbs development in ≥ 50% of the treated embryos when added up to 30 hpf (Fig. 3B1), while removing PFOA ceases to rescue normal development in ≥ 50% of the embryos at and after 21 hpf (Fig. 3B2). This broad window of effect from 21 to 30 hpf encompasses the morphological transition from late gastrula to pluteus stage, including the interval for secondary skeletal patterning (Fig. 3A) (Hogan et al., 2020; Piacentino et al., 2015; Piacentino, Zuch, et al., 2016). Most of the abnormalities observed in PFOA-treated embryos are broad skeletal patterning defects, with relatively few DV defects, consistent with the temporal period of action for PFOA (Fig. 3B1-B2). These results suggest that PFOA’s sufficiency for disrupting development during this interval reflects a late, potentially direct impact on pathways relevant to skeletal patterning. In contrast, Gen X exposure perturbed development when added up to 7 hpf (Fig. 3D1) and failed to rescue development when removed at 13 hpf (Fig. 3D2) when considering a 50% effect threshold. Unlike PFOA, most of the perturbations observed in Gen X-treated embryos are partial DV defects (Fig. 3D1-D2).The earlier window of effect from 7-13 hpf corresponds very well with DV specification (Bradham & McClay, 2006; Hardin et al., 1992; Piacentino et al., 2015). Overall, these data indicate that Gen X has an early, potentially direct effect on ectodermal DV specification that indirectly perturbs skeletal patterning, while PFOA has a later, possibly direct effect on skeletal patterning.

### PFOA and Gen X selectively perturb expression of ectodermal patterning cues and PMC genes

Sea urchin skeletal patterning cues include ectodermal VEGF, Wnt5, and the TGFß signal Univin, with the corresponding receptors expressed by the PMCs (VEGFR) or globally (Alk4/5/7) (Adomako-Ankomah & Ettensohn, 2013; Duloquin et al., 2007; Mcintyre et al., 2013; Piacentino et al., 2015; Piacentino, Zuch, et al., 2016; Thomas et al., 2023); the Wnt5 receptor is not known. The initially homogenous group of PMCs diversifies in response to these cues, with genes including *jun* and *pks2* expressed by spatially discrete PMC subsets (Adomako-Ankomah & Ettensohn, 2013; Hawkins et al., 2023; Sun & Ettensohn, 2014; Zuch & Bradham, 2019). The spicule matrix (SM) genes, including *msp130*, *sm30b*, and *sm50*, are also expressed by PMC subsets and encode proteins that are incorporated into the biomineral skeletal spicules (Benson et al., 1983; Lyons et al., 2014; Mann et al., 2010; Oliveri et al., 2002, 2003, 2008). We next tested how the expression of these genes was affected by PFAS exposure using HCR-FISH (Fig. 4-5).

**Figure 4:**
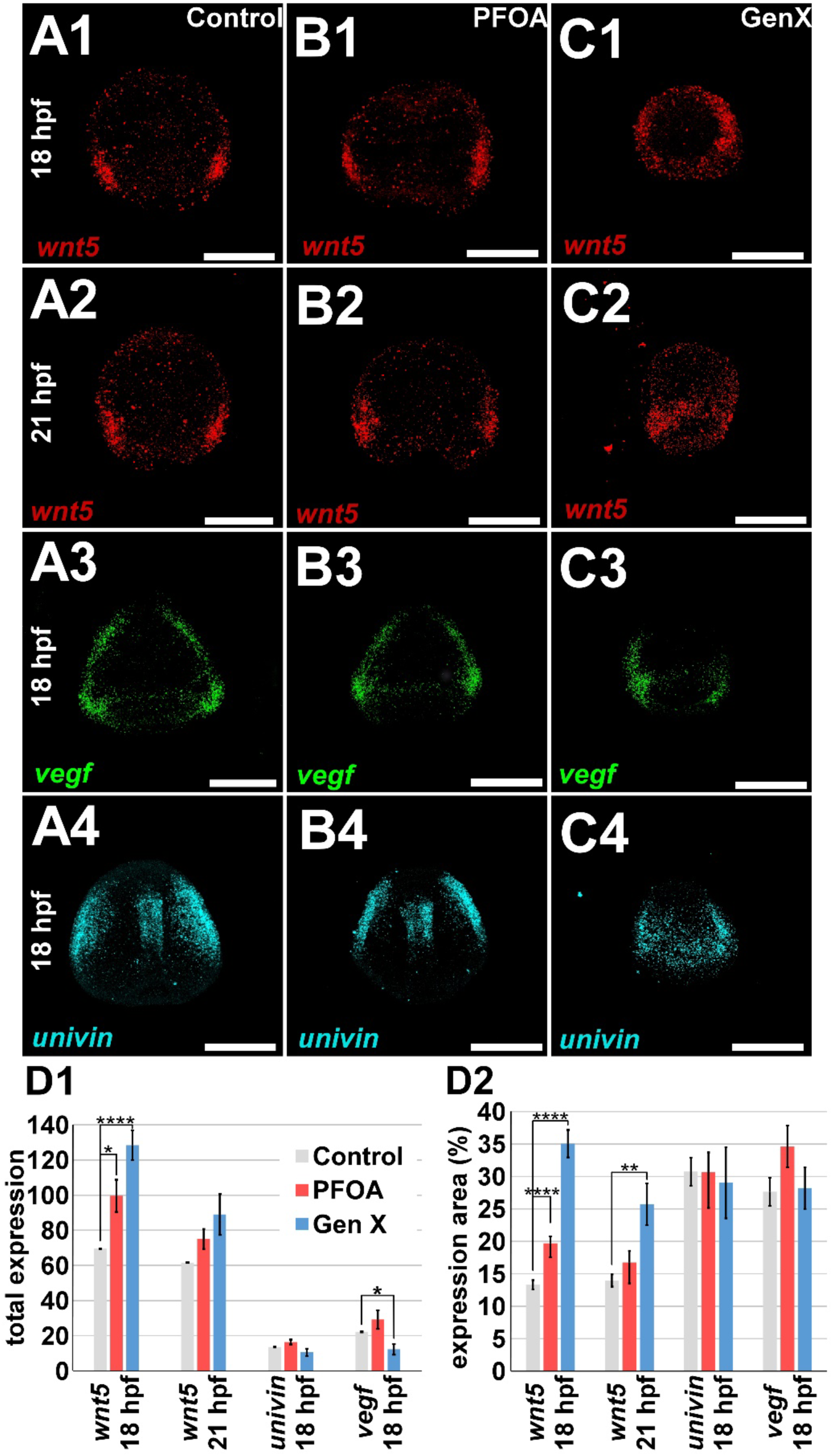
PFOA and Gen X are sufficient to perturb ectodermal patterning cue gene expression. **A-C** Control (A), PFOA-(300 μM, B) and Gen X-(250 μM, C) treated embryos were subject to HCR FISH for Lv-*wnt5* at 18 (1) and 21 (2) hpf, Lv-*vegfr* at 18 hpf (3) and Lv-*univin* at 18 hpf (4). **D.** Total expression (1) and the normalized area of expression (2) for each gene is plotted as the average ± s.e.m. * p < 0.05, ** p< 0.005, **** p < 0.00005; otherwise, insignificant (*t*-tests). See also Fig. S4.

**Figure 5:**
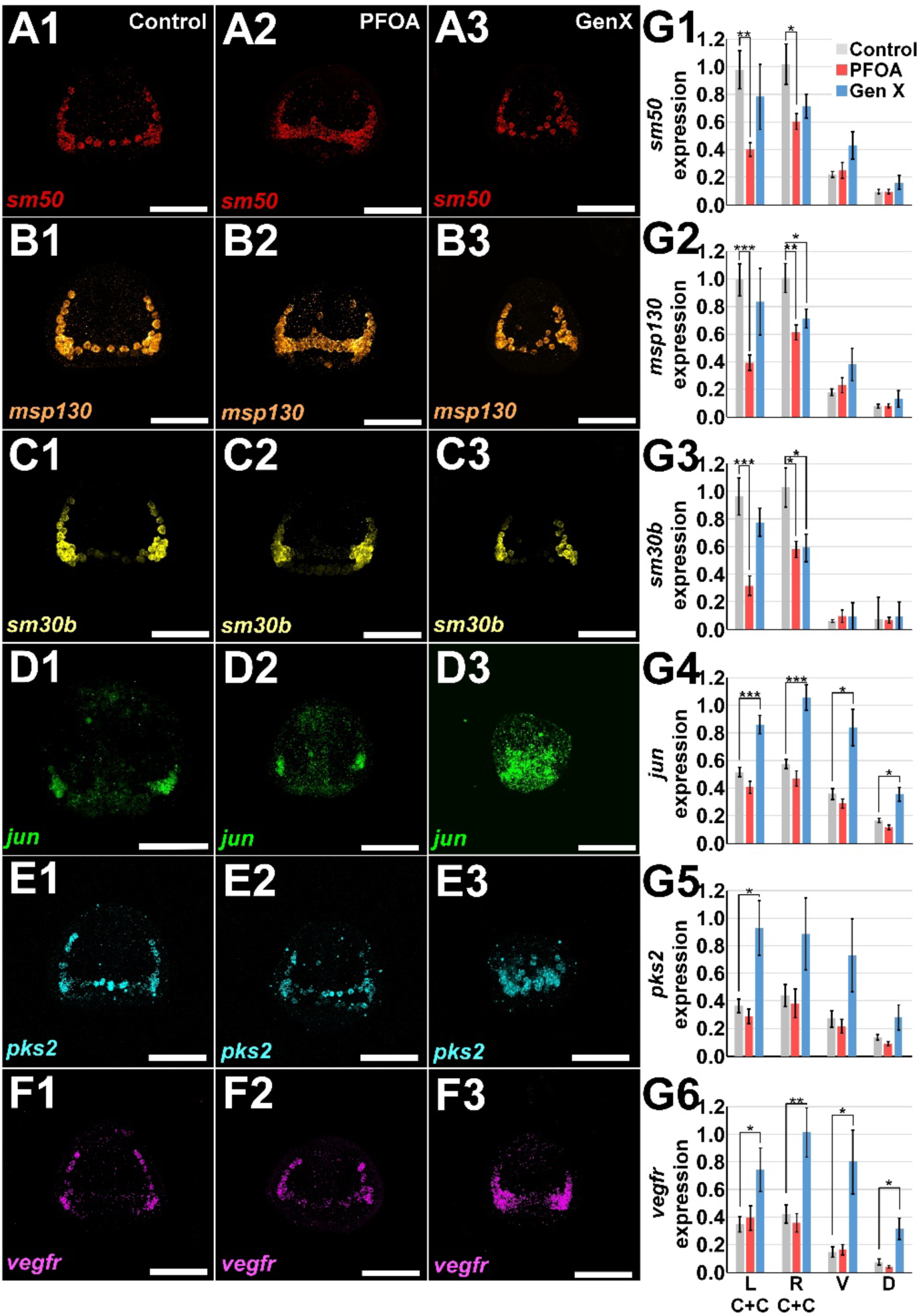
PFOA and Gen X are sufficient to perturb expression of PMC subset genes at 18 hpf. **A-F** Control (1), PFOA-(300 μM, 2) and Gen X-(250 μM, 3) treated embryos were subject to HCR FISH for Lv-*sm50* (A), Lv-*msp130* (B), Lv-*sm30b* (C), Lv-*jun* (D), Lv-*pks2* (E), and Lv-*vegfr* (F) at 18 hpf. **G**. The total expression (AFU x100) for each gene is plotted for the indicated spatial regions in the PMC pattern as the average ± s.e.m. * p < 0.05, ** p< 0.005, *** p< 0.0005; otherwise, insignificant (*t*-tests). L C+C, left clusters and cords; R C+C, right clusters and cords; V, ventral ring; D, dorsal ring. See also Fig. S5.

The patterning cue *wnt5* is expressed in the ventrolateral ectoderm at 18 and 21 hpf (Fig. 4A1-2) as expected (Mcintyre et al., 2013; Thomas et al., 2023); similarly, *vegf* and *univin* were normally expressed in ventrolateral or lateral ectodermal domains at 18 hpf (Adomako-Ankomah & Ettensohn, 2013; Duloquin et al., 2007; Piacentino et al., 2015; Piacentino, Zuch, et al., 2016) (Fig. 4A3-4). PFOA exposure produced only subtle effects on the expression of these patterning cues, while the impacts of Gen X treatment were more dramatic (Fig. 4B-C, S4). Quantification shows that neither *vegf* nor *univin* expression areas were affected by PFOA or Gen X exposure, nor was *univin* level at 18 hpf (Fig. 4D). Interestingly, Gen X-treated embryos frequently exhibited a significant unilateral reduction of *vegf* expression level (Fig. 4C3, D1). In contrast, *wnt5* level and expression area were significantly increased at 18 hpf by both PFAS, with Gen X exerting a more dramatic effect (Fig. 4A1-A2, B1-B2, C1-C2, D). At 21 hpf, PFOA- and Gen X-exposed embryos recovered normal *wnt5* expression level, while Gen X-exposed embryos were unable to laterally restrict the *wnt5* expression area (Fig. 4B1-2, C1-2, D1-2). Gen X exposure similarly spatially disrupted *univin* expression without changing its total expression area at 18 hpf (Fig. 4C4, D2); *univin* expression levels were reduced at 21 hpf by Gen X (Fig. S4). Thus, while exposure to either PFAS perturbed ectodermal patterning cue expression, Gen X produced a more severe and long-lasting spatial effect.

Because ectodermal cues impact PMCs, we next examined gene expression in that lineage. In control embryos, *sm50* is expressed by the ventral PMCs and the cords (Fig. 5A1, G1); *msp130* is most highly expressed in the PMC clusters and more weakly marks the cords and ventral part of the posterior PMC ring (Fig. 5B1, G2); *sm30b* expression is strongest in the PMC clusters (Fig. 5C1, G3). In PFOA-treated embryos at 18 hpf, these genes are each expressed at significantly reduced levels in the clusters and cords (Fig. 5A2, B2, C2, G1-3). Gen X mildly reduced right side *msp130* levels without impacting expression in the left side or either part of the ring at 18 hpf (Fig. 5A3, B3, C3, G1-3). Since biomineralization appears unaffected by either PFAS, these results imply that the perturbations of these spicule matrix-associated genes induced by PFAS exposure were not sufficient to disrupt biomineral formation.

In controls at 18 hpf, *jun* was expressed in PMC clusters, while *pks2* and *vegfr* exhibited a more complex pattern as expected (Fig. 5D1, E1, F1) (Adomako-Ankomah & Ettensohn, 2013; Duloquin et al., 2007; Hogan et al., 2020; Piacentino, Chung, et al., 2016; Sun & Ettensohn, 2014; Thomas et al., 2023; Zuch, D. T., & Bradham, 2019). The expression of both *jun* and *pks2* appears control-like in PFOA-treated embryos at 18 hpf (Fig. 5D-E, G4-5). In contrast, both *jun* and *pks2* exhibit abnormal, expanded spatial expression in Gen X-treated embryos at 18 hpf that is consistent with delayed development since at earlier stages, both *jun* and *pks2* are more broadly expressed within the PMC population, and, for *jun*, in the ectoderm as well (Hawkins et al., 2023; Rodríguez-Sastre et al., 2023; Thomas et al., 2023; Zuch & Bradham, 2019). Both genes are also upregulated in Gen X treated embryos compared to controls at 18 hpf, with *jun* being significantly overexpressed in each spatial subregion (Fig. 5G4) and *pks2* overexpressed in most (Fig 5G5). The lack of spatial restriction of *pks2* in Gen X-exposed embryos may be indicative of a reduction in LOX signaling (Hawkins et al., 2023). Gen X expanded *vegfr* expression throughout the PMCs at 18 hpf in a pattern that is consistent with a developmental delay but not with a strong DV perturbation (Duloquin et al., 2007). In contrast, *vegfr* expression was normal with PFOA at 18 hpf (Fig. 5F, G6). Together, these results show that PFOA has a stronger effect on spicule matrix protein-encoding gene expression, while Gen X instead effects expression of signaling components within PMCs during their patterning.

### PFOA and Gen X perturb DV spatial territories and ectodermal differentiation

To further investigate the impacts of PFOA and Gen X on DV specification, the spatial expression of DV marker genes was next assessed using HCR-FISH (Fig. 6A-C). We measured the expression of Lv-*chordin*, a target of Nodal signaling and a ventral marker, and Lv-*irxa*, a target of BMP2/4 signaling and a dorsal marker (Bradham et al., 2009; Su et al., 2009) (Fig. 6A-C). The ciliary band (CB), a morphologically distinct structure positioned at the dorsal-ventral (DV) boundary and comprised of apically constricted ciliated cells and neurons, is restricted to the space between the signaling domains of these two ligands. Perturbations to DV specification result in spatially aberrant CB patterning or restriction (Bradham et al., 2009; Yaguchi et al., 2010). In PFOA-treated embryos, the ventral domain is contracted, while the dorsal domain is correspondingly expanded, both in a small but statistically significant manner (Fig. 6B, D). This unexpected and surprising outcome indicates that PFOA treatment mildly impacts DV specification despite the evidence herein (see Fig. 3A-B) that PFOA has no effect during the DV specification interval from 7-13 hpf (hatched blastula to early gastrula stages) in Lv (Bradham & McClay, 2006; Hardin et al., 1992; Piacentino et al., 2015). These findings suggest that some spatial DV specification plasticity persists between 18-21 hpf in Lv, corresponding to late gastrula and prism stages, when PFOA exerts its effects.

**Figure 6.**
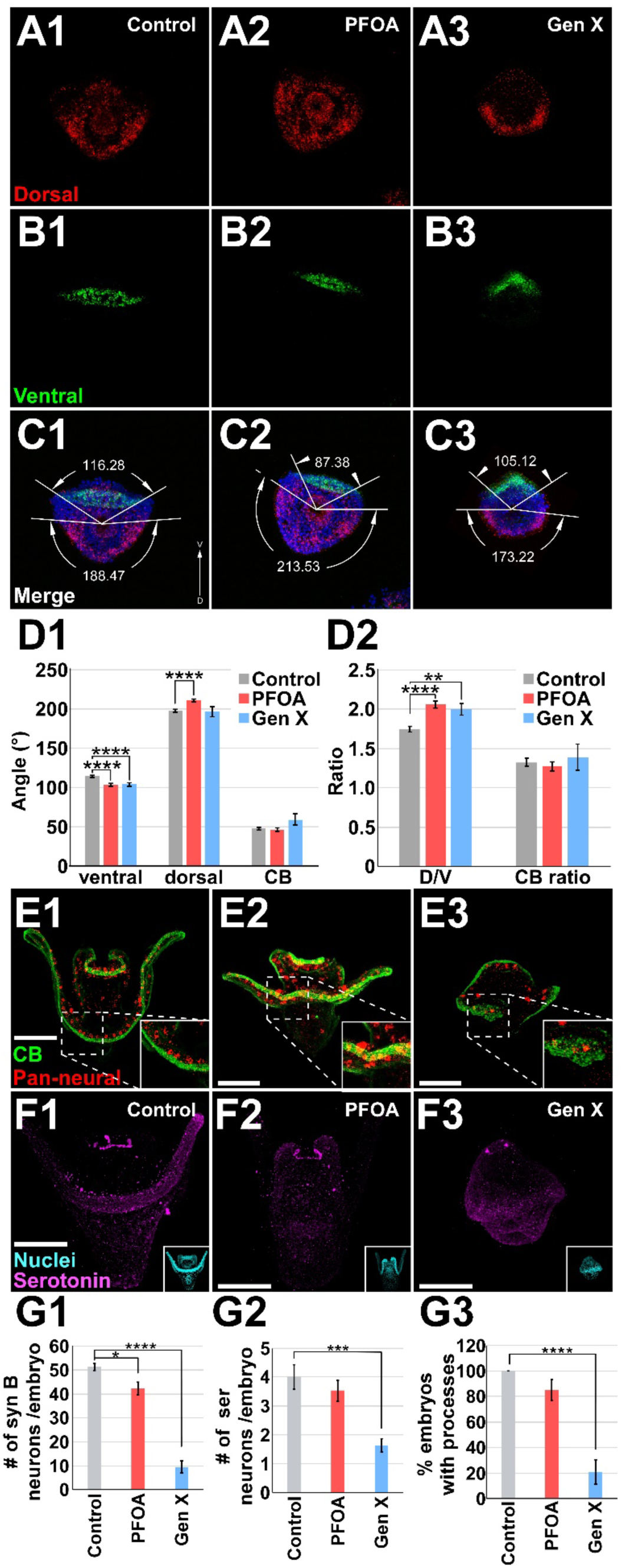
PFOA and Gen X are sufficient to perturb ectodermal DV specification and neural development. **A-C** Control embryos (1) and embryos treated with PFOA (300 μM, 2) or Gen X (250 μM, 3) were subject to HCR FISH for the dorsal marker Lv-*irxA* (red, A) and ventral marker Lv-*chordin* (green, B) at 18 hpf. Merged images that include Hoechst staining (blue, C) were used to measure the radial extent of Chd and IrxA expression domains as shown. **D**. The angular spatial extent of the ventral territory, dorsal territory, and the ciliary band (CB) are plotted as the average ± s.e.m. (1); the ratio of the dorsal and ventral expression levels and the ratio of the ciliary band angle to 360⁰ (2); ** p <0.005, *** p < 0.0005, **** p < 0.00005, otherwise not signficant (*t*-tests); n = 30 across 4 biological replicates. **E, F.** Control embryos (1) and embryos treated with PFOA (2) or Gen X (3) as above were subjected to HCR-FISH for the pan-neural marker synaptotagmin B (red) and immunolabeling for the ciliary band (green); insets show the extent of CB restriction at 48 hpf (E). Similar embryos were subjected to immunolabeling for serotonergic neurons (magenta, F), with corresponding Hoechst images inset. **G.** The number of synaptotagmin B neurons (G1), serotonergic neurons (G2), and number of embryos with evident processes extended by serotonergic neurons (G3) are plotted as average ± s.e.m; * p < 0.05, *** p< 0.0005, **** p < 0.00005; otherwise, insignificant. (*t*-tests); n = 11-20 per condition.

In contrast, Gen X-treated embryos show significant reduction to the ventral but not dorsal domain size. Instead, the CB, calculated as the space between the ventral and dorsal regions, is correspondingly enlarged but in a highly variable manner that does not reach significance (Fig. 6C, D). These results match the active window for Gen X that corresponds with DV specification (see Fig. 3C-D). Both treatments have a similar impact on the DV ratio; the fractional angle of the CB shows how the dorsal ratio increase relies on CB expansion with Gen X but not PFOA exposure (Fig. 6D2), implying that Gen X might directly impact the DV-specifying TGFß signals that control CB width, whereas PFOA appears to effect CB positioning, shifting it slightly ventrally without changing its size. Those differences in action agree well with the different temporal intervals of action for each chemical.

To test whether PFAS-treated embryos exhibit abnormal ectodermal differentiation, we visualized neurons and the ciliary band using HCR-FISH and immunolabeling (Fig. 6E-F). Neurons are derived from the ectoderm, and most neurons in sea urchin larvae are found in the ciliary band (Bisgrove, B. W., & Burke, 1986; Bradham et al., 2009; Burke R. D., 1978; Yaguchi et al., 2006), while the serotonergic neurons are localized in the apical part of the band (Yaguchi et al., 2006). We performed HCR-FISH to detect synaptogagmin B (synB), a pan-neural marker (Burke et al., 2006) and found that both PFOA- and Gen X-treated embryos have significantly reduced neuronal numbers overall (Fig. 6G1). This agrees with reduced neural specification in dorsalized embryos, especially in *L. variegatus* (Bradham et al., 2009; Yaguchi et al., 2006). In Gen X-treated embryos, the CB exhibits variable, localized abnormalities and discontinuities (Fig. 6E3 and inset), agreeing with results from earlier measures (Fig. 6A-D). In contrast, the CB in PFOA-treated embryos appears normal (Fig. 6E2 and inset). This lack of effect of PFOA on the CB also correlates with findings from earlier measures (Fig. 6A-D). Serotonergic (ser) neurons were detected by immunolabeling (Fig. 6F). PFOA-treated embryos exhibited no significant change in either neuron number or presence of processes for ser neurons; however, Gen X-treated embryos exhibited significant reductions in both ser neuron numbers and their processes (Fig. 6G2-3). These results show that PFAS exposure results in mild DV specification defects and, with Gen X especially, in neuronal losses.

### PFOA and Gen X are sufficient to perturb flow fields generated by larvae

The conspicuous cilia extended by cells in the CB drive the swimming behavior of the embryo. In cultures treated with PFOA, we observed altered swimming behavior with embryos floating at the surface of the water to form rafts, rather than swimming up and down in the water column (Movies S1-2). Gen X-treated larvae exhibit swimming behavior that is similar to controls aside from being slower (Movie S3). To test whether PFAS treatment perturbs cilia movement, we confined live embryos under coverslips, then continuously visualized particles in the media to quantitate the fluid flows using established particle image velocimetry (PIV) methods for data analysis (Stamhuis and Thielicke, 2014) (Fig. 7) (see Methods). Control 48 hpf larvae typically exhibit a flow pattern that consists of six vortices (Fig. 7A1). An obvious effect of the cilia movement and resultant vortices, particularly between the long (aboral) rods, is the generation of flow toward the larval mouth (Fig. 7B1, C1, D1); this is a novel observation. It is likely that the combined effects of the externally positioned vortices drives ventral-forward swimming.

**Figure 7.**
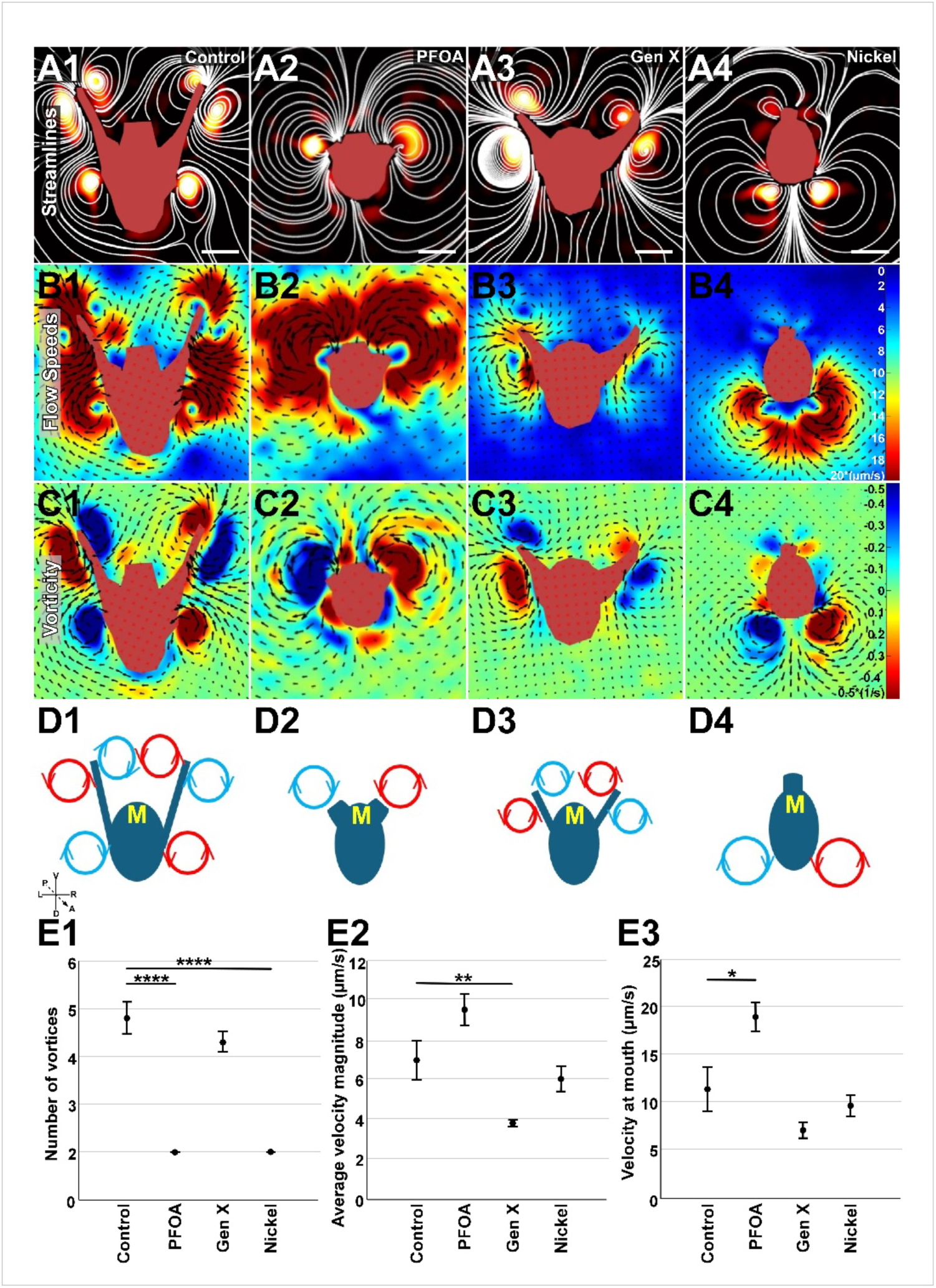
PFOA and Gen X are sufficient to perturb cilia movements and swimming behaviors. **A-D**. Exemplar control (1), PFOA-treated (300 μM, 2), Gen X-treated (100 μM, 3), and nickel-treated (0.2 μM, 4) embryos that were subjected to Particle Image Velocimetry (PIV) are shown as 3 second averaged snapshots with flow streamlines (white lines, A), and with vortex locator heatmaps that show flow speeds (B) or vorticity (C) with the embryos masked in red and with LUTs inset; the detected vortex locations and directions are schematized (D) with the mouth indicated (“M”). **E**. The average number of vortices generated (1), the average velocity magnitude around the larvae (2), and the average velocity magnitude at the mouth (3) are plotted as the average over 3 seconds ± s.e.m; * p < 0.05, ** p < 0.005, ****, p < 0.00005; otherwise, insignificant (*t* tests); n = 10. See also Fig. S5 and Movies S1-3.

When treated with PFOA, the resulting larvae exhibit an overall reduction in the number of vortices (Fig. 7A2, E1), particularly along the body, with bilateral flows only at the ends of each shortened larval arm (Fig. 7A2). While the average fluid velocity is increased in PFOA compared to controls (Fig. 7B1-2, C1-2), there is more flow into the mouth and less flow along the body (Fig. 7E2-3). These changes might explain the reduced swimming activity observed with PFOA (Movie S1-2). However, Gen X treatment resulted in distinct effects compared to PFOA. Gen X treatment typically reduced the number of vortices to four, with similar positioning to the vortices on the inner and outer faces of the larval arms as was observed in controls (Fig. 7A3, E1). However, strikingly, the fluid flows are strongly reduced with Gen X both around the larvae and at the mouth (Fig. 7B3, C3, D3, E2-3), agreeing with the loss of neurons previously observed with Gen X treatment (Fig. 6E-G). Because both Gen X and PFOA produced mild ventral contraction, we also evaluated Nickel-treated embryos, which exhibit ventralization (Hardin et al., 1992) to evaluate how DV perturbation impacts fluid flows. Nickel-treated larvae were used at 48 hpf and exhibit a typical morphology for this developmental stage (Fig. S5D). The flows around Ni-treated larvae have a very different profile, with vortices only at the posterior end of the body (Fig. 6A4). This location corresponds to the repositioned CB in ventralized embryos (Bradham et al., 2009; Duboc et al., 2004; Yaguchi et al., 2006).

Unexpectedly, most PFOA-treated embryos (8/10) exhibited left or right-dominated flow patterns (Fig. S5A). This LR asymmetry also occurred with Ni (Fig. S5B). Curiously, these LR asymmetries were randomly distributed among both groups, and were also present, with less variation, with Gen X and in controls (Fig. S5C). These results suggest that PFAS exposure perturbs the neural functions that regulate cilia movements, matching their effects on neural specification (Fig. 6E-G). In keeping with their impacts on neural development, Gen X produces larger deficits in ciliary function than PFOA.

## DISCUSSION

In this study, we characterize and compare the effects of the PFAS chemicals PFOA and Gen X on *L. variegatus* development. We demonstrate that each chemical is sufficient to perturb *L. variegatus* development, with impacts on endomesoderm specification, the DV axis, skeletal patterning, and neural development and function. We establish that PFOA and Gen X act during distinct temporal windows during development, with Gen X acting earlier than PFOA, suggesting that they differ in their mechanisms of action. Endomesoderm specification is strongly delayed or inhibited by Gen X. The ventral ectoderm is mildly contracted by each chemical, via dorsal expansion with PFOA versus ciliary band expansion with Gen X. Each treatment perturbs skeletal patterning, with differing impacts on the expression of patterning cues in the ectoderm and probable cue-response genes in the PMCs. The timing results and specific skeletal abnormalities combined with the impacts of PFAS exposure on ectodermal and PMC gene expression patterns suggest that PFOA more directly perturbs skeletal patterning, while Gen X acts earlier, albeit mildly, on the DV specification pathway; these effects result in distinct skeletal patterning defects. Gen X exposure dramatically perturbs neural specification and function, which is much less affected by PFOA. These overall findings are summarized in Figure 8.

**Figure 8.**
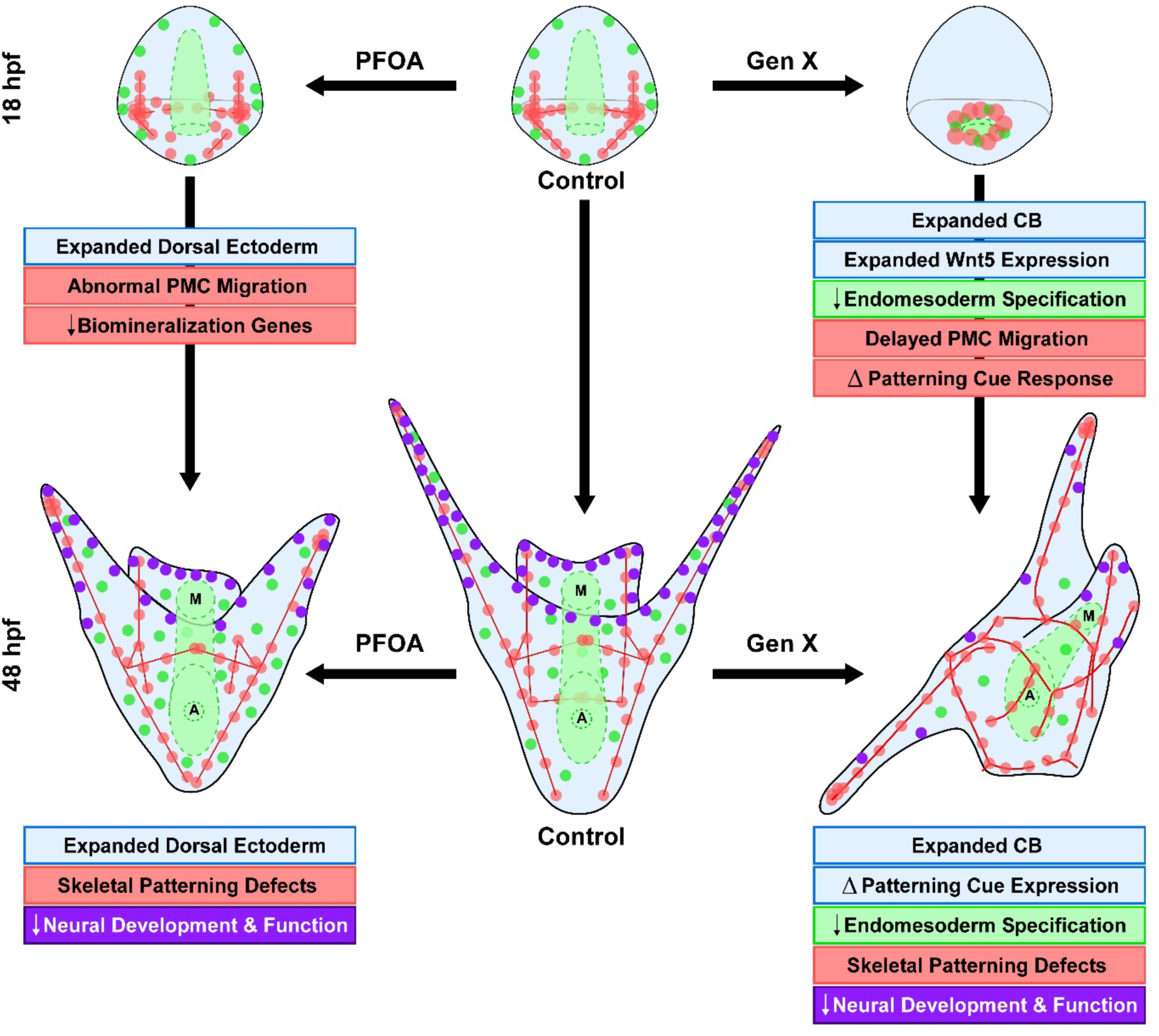
A summary of the perturbing effects of PFOA and Gen X on *Lytechinus variegatus* embryos and larvae is shown at three time points. Changes in the ectoderm (blue), endomesoderm (green), PMCs and skeleton (red), neurons (purple), morphogenesis, and gene expression are depicted for controls (center) and embryos treated with PFOA (left) or Gen X (right). M, mouth; A, anus.

To our knowledge, this is the first study that assesses the effects of PFAS exposure on developing sea urchins. Prior work has instead focused on the effects of PFAS on adult sea urchins (Hamed et al., 2024; Hayman et al., 2021) or on embryos reared from exposed parents; interestingly, such embryos also exhibit patterning defects (Savoca et al., 2023). Contrary to other studies which reported reduced developmental toxicity with Gen X compared to legacy PFAS (Degitz et al., 2024; Gaballah et al., 2020; Gebreab et al., 2020; Satbhai et al., 2022), our study indicates that Gen X is more teratogenic than PFOA, in part because it acts earlier in development, impacting axial and germ layer specification, unlike PFOA.

PFAS exposure also interfered with endomesoderm differentiation. This effect was much more pronounced and prolonged with Gen X, while PFOA exposure appears to only mildly delay gut morphogenesis with no evident impact on SMC specification.

Because PFOA acts later in development, after the body axes and germ layers are thought to be specified, perhaps that timing explains its comparatively mild effect on the gut. In contrast, Gen X acts much earlier, during secondary axis specification, when it could potentially interfere with endomesoderm specification more directly. Even so, Gen X-treated embryos do eventually gastrulate, and appear to have complete through-guts by the larval stage.

The strong inhibition of neural specification and function with Gen X reported herein agrees with other studies that show an increase in expression of neurotoxicity-associated gene transcripts after treatment with Gen X as well as an increase in repressive histone marks associated with neurodegeneration (Wu et al., 2023). Thus, the neurotoxic effects of Gen X on developing embryos observed herein may be generalizable.

Skeletal patterning defects are the second-most common birth defect world-wide, yet their etiology remains mechanistically unclear (Arth et al., 2017; Cornel, 2000; Eide et al., 2006; Hinton et al., 2017; Mai et al., 2019; Petterson et al., 2007). Interestingly the pathways involved in sea urchin skeletal patterning are often involved in human skeletal development (Cottrell et al., 2013; Garcia et al., 1996; Hästbacka et al., 1994; Ivanova et al., 2022; Kere, 2006; Mcintyre et al., 2013; Oh et al., 2023; Piacentino et al., 2015, 2016; Rys et al., 2016; Thomas et al., 2023; Traianedes et al., 1998; Wu et al., 2016; Xu et al., 2023). In sea urchin embryos, Gen X exposure perturbs the ectodermal skeletal patterning cues Wnt5, VEGF, and Univin, while PFOA exposure more transiently impacted Wnt5 without affecting the other two cues. PFAS exposure also impacted the cellular behavior and gene expression dynamics of the PMCs during skeletogenesis. Gen X-treated embryos exhibited spatial expression patterns typical of earlier stages for *jun* and *pks2*, possibly indicating a lack or delay of patterning cue reception by the PMCs and consistent with Gen X-mediated perturbation of cue expression. In contrast, PFOA reduced the expression of skeletal differentiation genes without impacting *jun* or *pks2*, underscoring the distinct mechanisms of action of these PFAS molecules. These findings agree with other studies showing that PFAS exposure is sufficient to perturb skeletal growth and development in both pubescent and embryonic mice (Feng et al., 2024; Koskela et al., 2016); thus, like the neuronal defects, the developmental skeletal patterning defects observed herein in echinoderm embryos and larvae may be generalizable to vertebrates and mammals.

Overall, these results identify both PFOA and Gen X as potent teratogens for echinoderms, with Gen X exhibiting more profoundly toxic effects, including impacts on the specification of endomesoderm, the secondary body axis, neural development and function, and pattern formation. While some of these effects may be generalizable, similar studies in frogs and zebrafish did not identify axial defects in embryos treated with PFOA or Gen X (Degitz et al., 2024; Gaballah et al., 2020). Degitz et al. observed developmental delay in *Xenopus laevis* embryos at a low dose of 15 μM for both PFOA and Gen X, but oddly not with doses comparable to those in this study. The authors speculate that the genetics of the breeding pairs used for the studies may have affected this outcome. Similarly, Gaballah et al. utilized doses at or below 80 μM for both PFOA and Gen X and observed no phenotypic defect in frogs or fish; however earlier studies utilizing PFOA in zebrafish found malformations of the head and tail at 2 μM that could be indicative of axial defects when treating closer to fertilization (Hagenaars et al., 2013; Jantzen et al., 2016).

The ecological significance of these findings is likely to grow as environmental concentrations of PFAS chemicals rise, particularly in view of the lack of regulation surrounding Gen X as compared to legacy PFAS like PFOA (perfluorooctanoic acid) and PFOS (perfluorooctanesulfonic acid) that are currently recognized by the USA and many other countries as hazardous substances and therefore banned (Langenbach & Wilson, 2021). Sea urchin embryos are recognized as sentinel marine animals (Arnberg et al., 2018; Cossa et al., 2024; Dorey et al., 2022; Duvane & Dupont, 2024; Moreno-García et al., 2022; Pinsino & Matranga, 2015; Przeslawski et al., 2015; Sarly et al., 2023; Sato et al., 2018) and as such, they provide an early warning of the potential negative consequences of the continual environmental accumulation of these “forever” chemicals to vertebrate and mammalian species, including humans.

The evidently distinct modes of action observed herein for PFOA and Gen X agree with other findings that also indicate differences in their toxicological mechanisms (Attema et al., 2022; Bangma et al., 2021; Blake et al., 2020; Wen et al., 2020; Yoo et al., 2023). The relatively high potency of Gen X agrees with studies employing a placental trophoblast model, which showed that the parent chemical for Gen X both accumulated intercellularly at lower concentrations than PFOA, and provoked stronger effects on gene expression (Bangma et al., 2021; Bischel et al., 2011). Thus, in Gen X-exposed mammals, developing embryos will experience a higher-than-systemic level of Gen X due to placental enrichment. Moreover, the placenta is highly sensitive to the adverse effects caused by Gen X which include atrophy, necrosis and congestion of the labyrinth, resulting in placental insufficiency and reduced nutrient and oxygen transfer to the fetus (Blake et al., 2020; Blake & Fenton, 2020; Conley et al., 2019; Lv et al., 2024), underlining our and others observations that the potency and danger of the effects of Gen X exceed those of already-banned PFOA, and calling for policy changes that also ban the use of Gen X.

Because these chemicals are so stable and persistent in the environment and within biological systems, they will continue to accumulate and their concentrations will continue to increase if strategies to remediate them are not actualized, even if their production were to cease. Thus, determining their effects on biological systems at relatively high doses is ecologically relevant; seeking methods for their removal is crucial for the prevention of widespread teratogenesis both environmentally and among humans.

## Materials and Methods

### Animals, Chemicals, and Drug Treatments

Adult *Lytechinus variegatus* sea urchins were obtained from Pelagic Corp (Sugarloaf Key, FL, USA), the Duke University Marine Lab (Beaufort, NC, USA) or from Jake Warner or Laura Salter (NC). Gametes were collected by injecting adult sea urchins with 0.5 M potassium chloride solution. High purity perfluorooctanoic acid (PFOA, CAS No. 335-67-1) and hexafluoropropylene oxide dimer acid (Gen X, CAS No. 13252-13-6) were obtained from Sigma-Aldrich (WI, USA) or Synquest Laboratories (FL, USA), respectively. Chemical stock solutions were frozen in single-use aliquots to avoid repeated freeze-thaw cycles. Embryos were cultured in artificial sea water (ASW) or ASW and PFAS. ASW was buffered with 1 µL saturated NaHCO_3_ solution per 1 mL ASW (Thomas et al., 2023). PFAS optimal doses were determined as 300 µM for PFOA and 250 µM for Gen X via dose-response experiments geared towards identifying the minimal dose that produced penetrant, reproducible, and non-lethal perturbations. Those doses were employed in each experiment except when stated otherwise. Some early experiments used a Gen X solution at 875 µM; however, based on subsequent optimization studies, we are skeptical that that dose was accurate. Chemicals were added at fertilization unless otherwise noted.

### Skeletal Imaging and Time Course Experiments

Sea urchin skeletal images were obtained via illumination with plane-polarized light to capture the birefringence of the skeletal crystal. Images of each embryo’s skeleton were taken in multiple focal planes, then assembled manually in Photoshop (Adobe; v 22.0.1) or Canvas X Draw (v 20 build 914) to visualize the complete skeleton in focus. For PFAS removal experiments, PFOA or Gen X was added at fertilization, then removed at various time points. For PFAS addition experiments, sea urchin cultures were fertilized in ASW, then PFOA or Gen X was added at various time points. For both experimental series, the embryos were developed to the pluteus larval stage (48 hpf), photographed, then analyzed from the images. Embryos were scored and counted in Canvas X Draw (Canvas GFX; v 7.0.2).

### Immunolabeling, Hybridization Chain Reaction Florescent In Situ Hybridization (HCR FISH), and confocal microscopy

For PMC immunostaining, embryos were fixed in 4% paraformaldehyde in ASW for one hour for compatibility with HCR-FISH; otherwise, fixation and staining were performed as described (Bradham et al., 2009; Gross et al., 2003). Primary antibodies include PMC-specific monoclonal 6a9 (used at 1:30 or 1:50; from Prof. Charles Ettensohn, Carnegie Mellon University, Pittsburgh, PA, USA), ciliary band-specific monoclonal 295 (undiluted; from Prof. David McClay, Duke University, Durham, NC, USA), and serotonergic neuron-specific polyclonal α-ser (1:1000, Sigma Aldrich). Secondary antibodies were used at 1:500 and include goat anti-mouse antibodies labeled with Alexa Fluor 488 (Thermo Fisher), Alexa Flour 546 (Thermo Fisher), or DyLight 405 (Jackson Laboratories), and Cy2-conjugated goat anti-rabbit (1:900; Jackson Laboratories). Hoechst nuclear stain (Sigma Aldrich) was added at 1:500 with secondary antibody for some experiments. For HCR-FISH, embryos were fixed in 4% paraformaldehyde in ASW for one hour. HCR-FISH was performed as described (Choi et al., 2016) using hairpins and probe sequences designed by Molecular Instruments, Inc. In some cases, embryos were subjected to HCR-FISH followed by immunolabeling. Confocal microscopy was performed with Nikon C2 or Olympus FV10i-DOC microscopes at 200X or 400X; the corresponding z-stack projections were produced with FIJI (v 2.1.0); all z-stack images are shown as full projections.

### DV, Neural Analysis, and SMC analysis

For dorsal-ventral HCR-FISH analyses, the extent of *chordin* and *irxa* spatial expression was measured radially from the center of the gut using the angle measurement tool in ACD Canvas, and the ciliary band size was calculated by subtracting the sum of *chordin* (ventral) and *irxa* (dorsal) angle measurements from 360° on a per-embryo basis (Thomas et al., 2023). Neuron counts per embryo were quantified manually by inspection of unprojected z-stack images of neural immunolabels or HCR-FISH images of pan-neural marker synaptotagmin B. SMCs were similarly counted from HCR-FISH images of *Lv-pks1* expression. Statistical significance was determined using unpaired two-tailed student’s *t*-tests for pair-wise comparisons.

### Ectodermal, PMC, and Endodermal Gene Expression Quantifications

Quantification of HCR-FISH signals for ectodermal gene expression was performed in FIJI using maximum projections of confocal z-stacks (Fig. S6C). Regions of interest (ROIs) were drawn around discrete areas of expression within the ectoderm for each gene, then the integrated density and area for each ROI was obtained. An additional ROI was drawn around the circumference of the embryo. The total gene expression was obtained by summing the integrated density for each ROI per embryo, and the total area of expression for each gene was calculated by summing the area for each of the ROIs per embryo, then dividing by the overall area of the embryo for normalization. Quantification of HCR-FISH signals for PMC gene expression was performed in FIJI using maximum projections of confocal z-stacks that were rotated in Napari to a uniform orientation to most accurately capture dorsal and ventral signal differences (Fig. S6A-B). ROIs for PMC genes were drawn to encompass PMCs in the left and right cords and clusters, and the ventral and dorsal portion of the PMC ring. Expression levels were defined as the integrated density within the ROI. Endodermal gene expression level and expression area were similarly obtained from ROIs drawn around the *endo16* expression area in maximum projected confocal z-stacks. Values were then averaged for each condition, and statistical significance was determined using unpaired two-tailed student’s *t*-tests for pair-wise comparisons.

### PIV and Flowtrace Analysis

To quantify the fluid flows and their patterns around control and chemically treated sea urchin larvae, live imaging experiments were performed under squeeze-confinement conditions. First, glass slides were prepared for imaging by attaching a pair of long rectangular strips of double-sided tape of 50 µm thickness (Nitto Inc.) side by side to serve as spacers. Next, a droplet (∼50 µl) of seawater containing the sea urchin larva was applied between the spacers. A second droplet of (∼ 10 µl) of a solution of 1 µm polystyrene microspheres (Polysciences, cat. no. 07310-15) was then added and gently mixed; their final dilution was around 1:100 (volume of beads to volume of sea water). A cover slip was then slowly introduced onto the glass slide, then attached to the spacers, resulting in a confinement chamber that gently trapped the 100 µm sea urchin larvae under 50 µm squeeze-confinement conditions. We then carried out live darkfield imaging to quantify the movement of the microspheres with a Zeiss Axio Imager M2 (an upright microscope) using a 10X objective to capture a rectangular field of view of 1.5 mm X 1.5 mm. We captured time-lapse images at 10 fps for 30 seconds using a high-speed camera (Hamamatsu ORCA-Fusion Gen-III sCMOS). The time-lapse images were post-processed using PIVlab in MATLAB (Stamhuis, E. and Thielicke, 2014) to quantify the larval fluid flows. To meet the objective of quantifying the flows surrounding the larvae, a mask was drawn over the larva to avoid any flow quantification artifacts inside the body due to noise. The set of images was pre-processed to remove background noise using high-pass filters prior to the PIV analysis step, which calculates the time-varying velocity vector fields. The velocity vector fields were post-processed to smoothen the fields and spatially interpolate vectors if necessary. Finally, the fluid flow parameter of interest (vorticity or velocity magnitude) was plotted along with the velocity vector fields. To obtain smooth fluid flow visualizations and quantification, the averaged mean of the parameter of interest over a time window of 30 frames (corresponding to a time scale of 3 seconds) was calculated.

## Acknowledgements

We would like to thank Professors Charles Ettensohn and David McClay for antibodies. We would also like to thank Todd Blute for equipment training and support.

## CRediT authorship contribution statement

Author contributions: **A.T. Lion**: methodology, investigation, formal analysis, validation, visualization, writing-original draft, writing-review and editing **S.M. Bodine**: investigation, formal analysis, validation, visualization, writing-review and editing **K.R. McCutcheon**: investigation, validation, writing-review and editing **M. Ghogale**: visualization, formal analysis, writing-review and editing **S. Chandragiri**: investigation, data curation, visualization, formal analysis, writing-review and editing **D. Abayawardena**: investigation, visualization, writing-review and editing **B. Shrestha**: investigation, data curation, visualization, formal analysis, writing-review and editing **A. Descoteaux**: investigation, visualization, writing-review and editing **K. Alvarez**: investigation, formal analysis, visualization **J. A. Balkman**: investigation, formal analysis, visualization **B. Cocke**: investigation, formal analysis, visualization **A.H. Wikramanayake**: supervision, writing-review and editing **J. Schlezinger**: resources, writing-review and editing **J.Y. Wong**: conceptualization, writing-review and editing **V.N. Prakash**: methodology, supervision, writing-review and editing **C.A. Bradham**: conceptualization, methodology, supervision, project administration, funding acquisition, visualization, formal analysis, writing review and editing

