## Supplemental Figures and Legends for "PFAS Compounds PFOA and Gen X are Teratogenic to Sea Urchin Embryos"

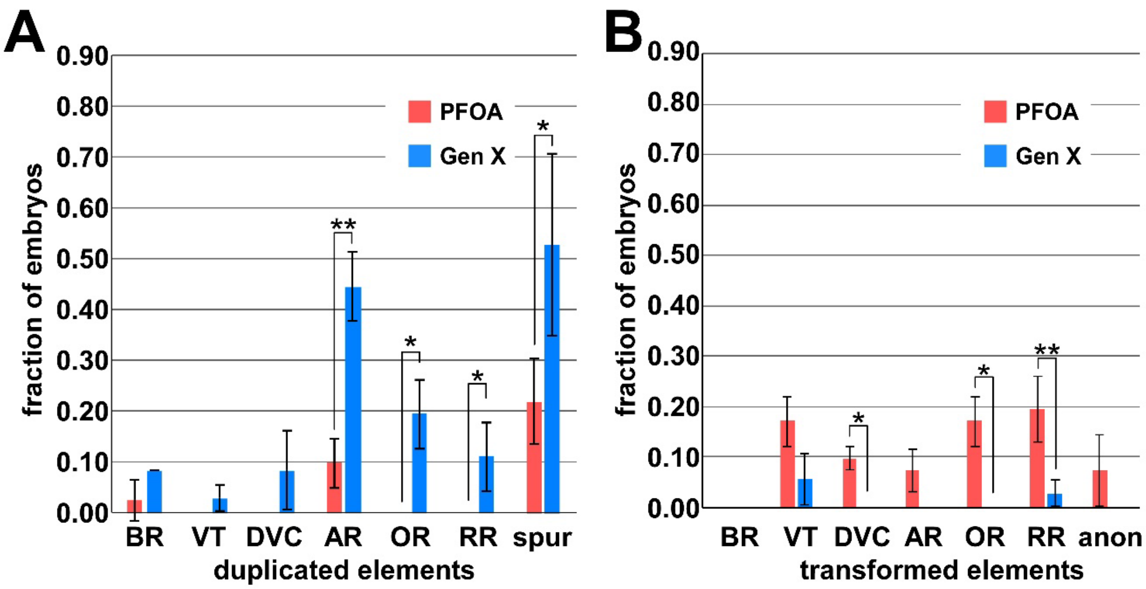


**Figure S1. Embryos treated with PFOA and Gen X have differing patterning defects.** Gen X-treated embryos exhibit significantly more duplicated elements than PFOA-treated embryos (**A**), whereas PFOA treated embryos have significantly more transformed elements than Gen X-treated embryos (**B**). See also Fig. 2.

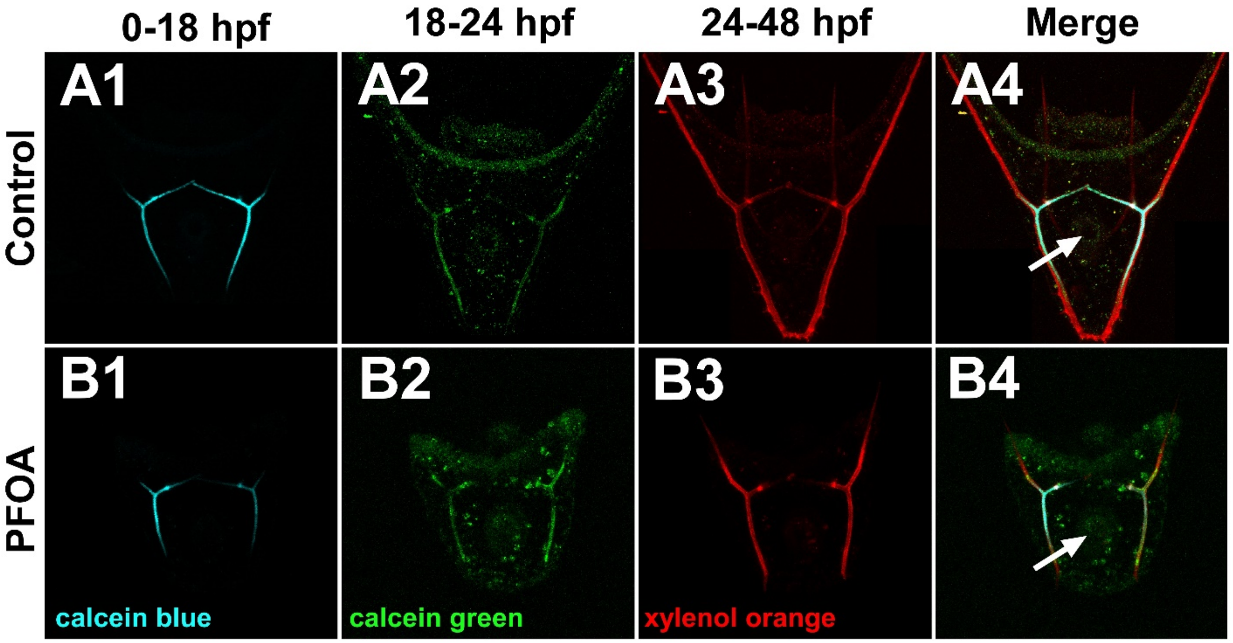


**Figure S2. Birefringence signals in the gut of PFOA-treated embryos does not contain calcium.** Calcium was visualized using three polychromes that were applied for the indicated temporal intervals (**1-3**) and merged (**4**) in control (**A**) and PFOA treated embryos (300 µM, **B**). In both cases, calcium is detected in the skeleton but not the larval gut (arrowheads). See also Fig. 2.

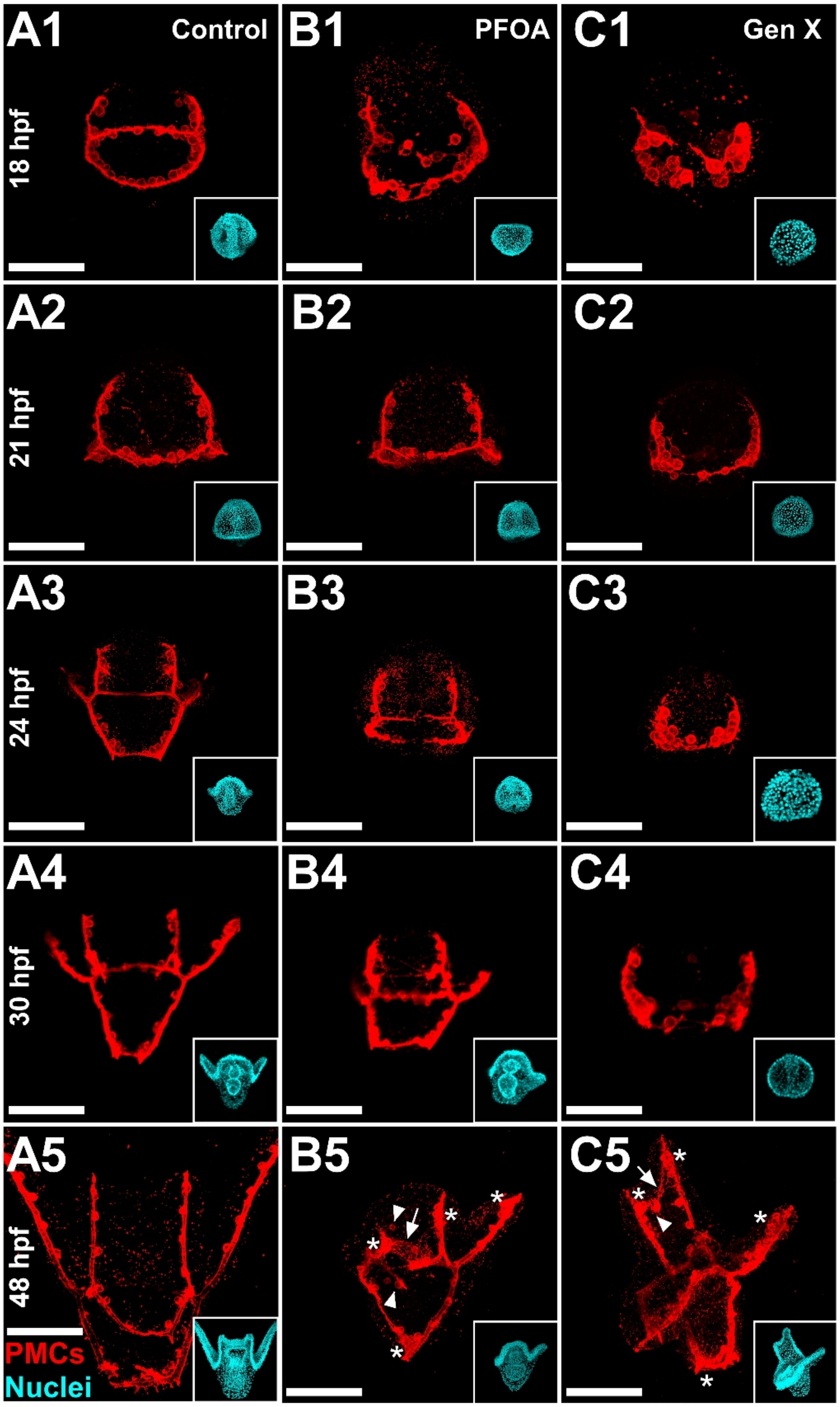


**Figure S3. PMC migration and positioning is perturbed by PFOA or Gen X treatment.** Time course of PMC migration in Control (**1**), PFOA (**2**), or Gen X (**3**) embryos at 18 (**A**), 21 (**B**), 24 (**C**), 30 (**D**), and 48 hpf (**E**). Some of these results are also shown in Figure 2.

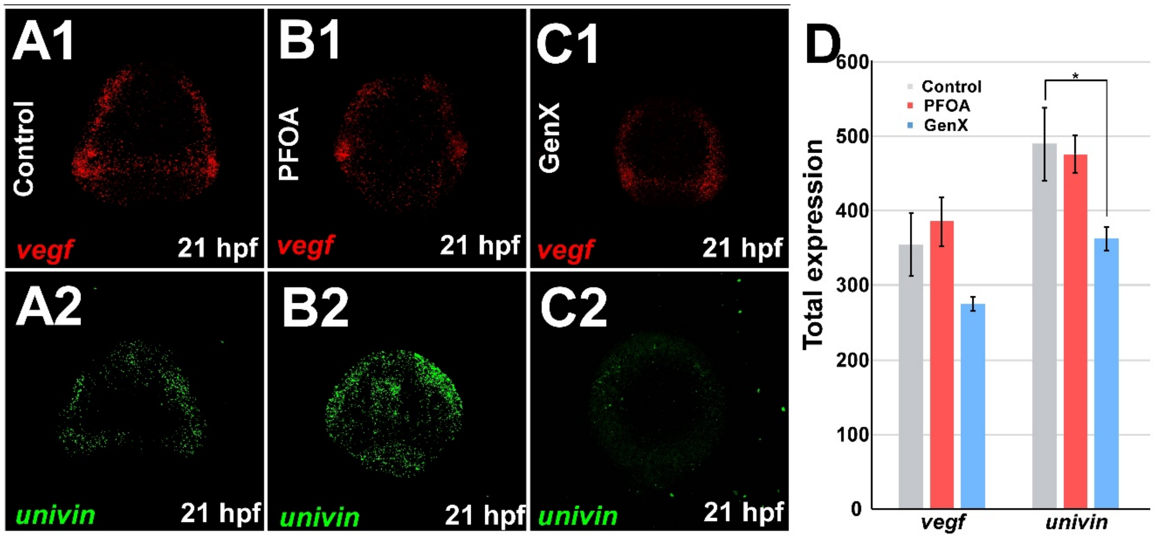


**Figure S4. PFAS perturb ectodermal patterning cue expression.** Control (**A**), PFOA- (300 µM, **B**) and Gen X- (250 µM, **C**) treated embryos were subject to HCR-FISH for Lv-*vegf* (**1**) or Lv-univin (**2**) at 21 hpf. **D**. The total expression for each gene is plotted as the average ± s.e.m., * $p < 0.05$ (*t*-tests). See also Fig. 4.

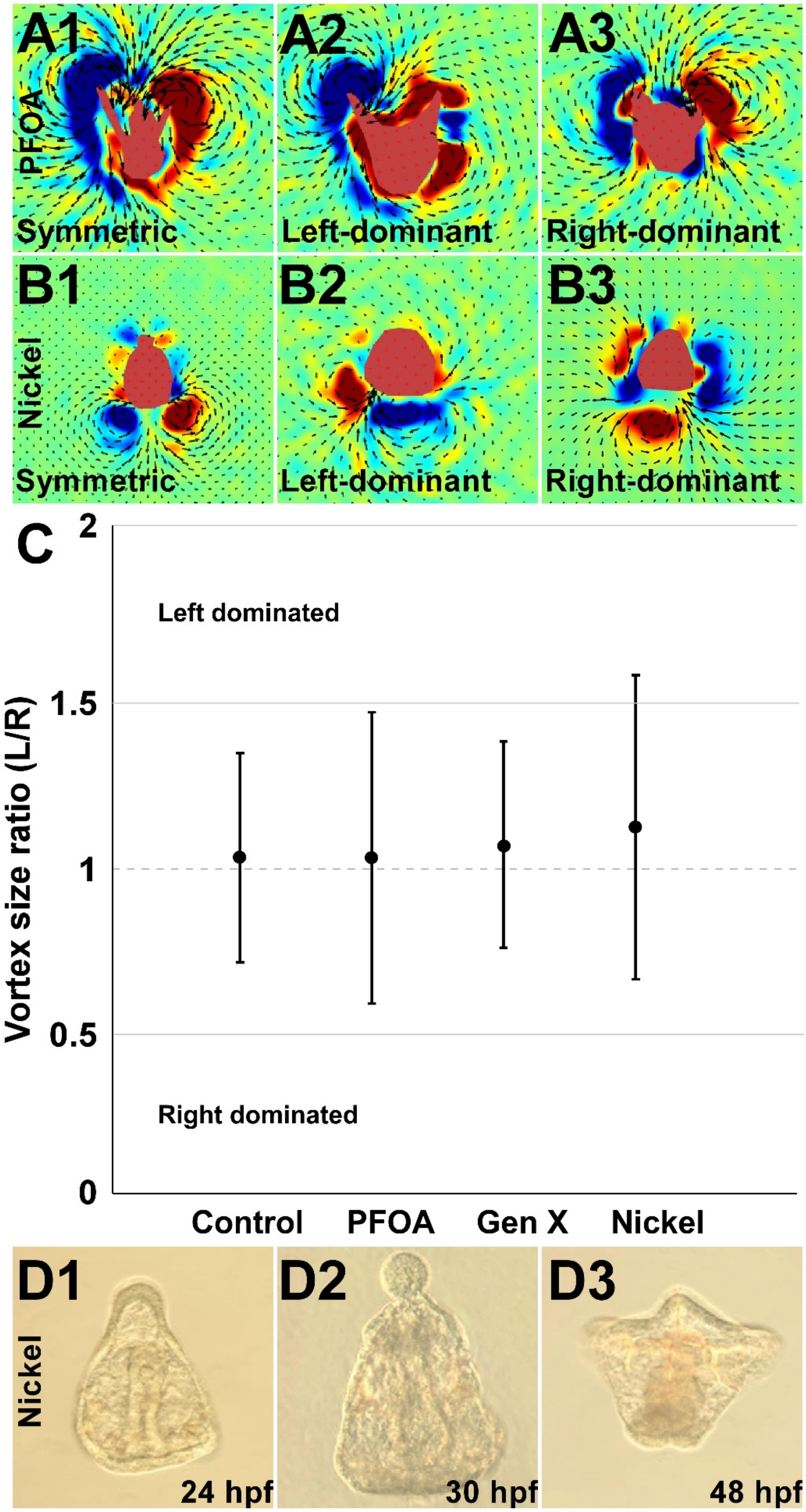


**Figure S5. Fluid flows around PFOA- and Nickel- treated embryos exhibit left-right asymmetry in vorticity, but not consistently on one side. A-B.** Exemplar PFOA- (300 µM, **A**) or Ni- (0.2 µM, **B**) treated embryos showing symmetric (**1**) or left-right asymmetric (**2-3**) vorticity. **C**. The left-right vortex size ratio is plotted as the average ± standard deviation for the indicated treatments. **D**. The morphology of Nickel-treated embryos is shown at the indicated time points as DIC images. See also Fig. 8

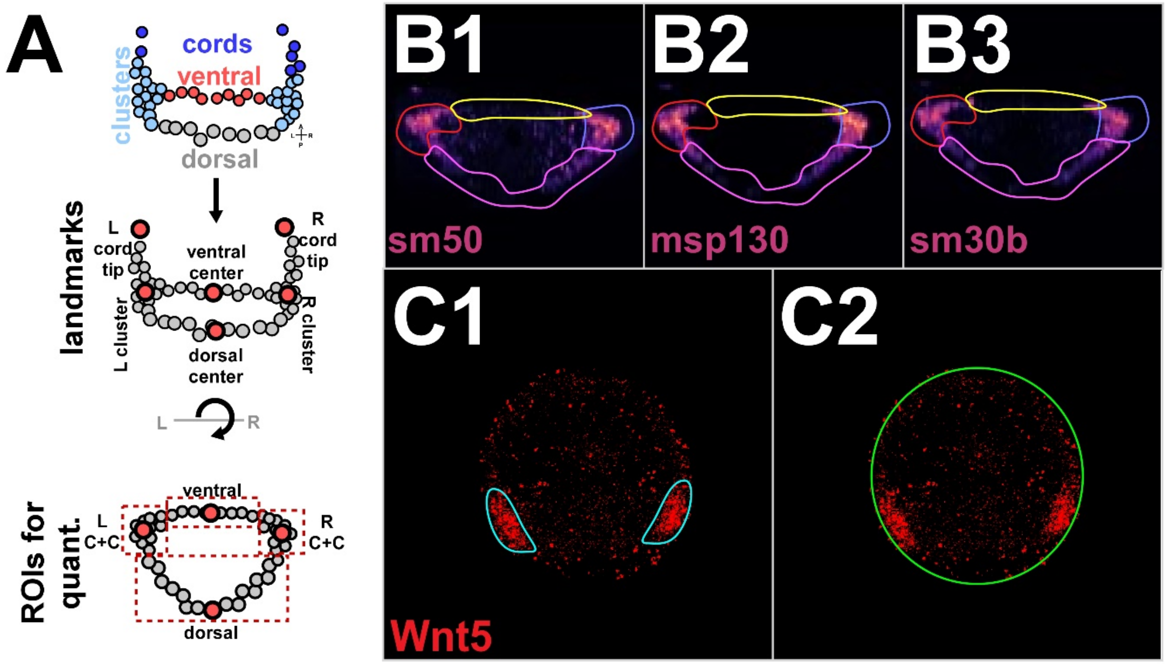


**Figure S6. Methodology for quantifying expression level and area from HCR-FISH images. A**. Embryos were subjected to HCR-FISH for a set of PMC genes and/or ectodermal patterning cues (see Fig. 5-6 and S5-6). Schematics depict confocal z-stack projections (top) that are landmarked as shown (middle), then rotated using Napari to a uniform vegetal view in which cord and cluster landmarks are aligned (bottom). ROIs or PMC subregions are schematized (bottom). **B.** For PMC genes, ROIs are drawn around the clusters and cords (left, red; right, blue), ventral ring (yellow) and dorsal ring (magenta). **C.** For ectodermal cues, ROIs were drawn that enclose the bilateral areas of expression (**1**), and the embryo (**2**). The gene expression area was normalized to the area of the embryo.
